# Best practices for perturbation MPRA—a computational evaluation framework of sequence design strategies

**DOI:** 10.1101/2023.09.27.559768

**Authors:** Jiayi Liu, Tal Ashuach, Fumitaka Inoue, Nadav Ahituv, Nir Yosef, Anat Kreimer

## Abstract

The advent of the perturbation-based massively parallel reporter assays (MPRAs) technique has enabled delineating of the roles of non-coding regulatory elements in orchestrating gene expression. However, computational efforts remain scant to evaluate and establish guidelines for sequence design strategies for perturbation MPRAs. Here, we propose a framework for evaluating and comparing various perturbation strategies for MPRA experiments. Under this framework, we benchmark three different perturbation approaches from the perspectives of alteration in motif-based profiles, consistency of MPRA outputs, and robustness of models that predict the activities of putative regulatory motifs. Although our analyses show similar while significant results in multiple metrics, the method of randomly shuffling nucleotides outperform the other two methods. Thus, we still recommend designing sequences by randomly shuffling the nucleotides of the perturbed site in perturbation-MPRA. The evaluation framework, together with the benchmarking findings in our work, creates a resource of computational pipelines and illustrates the promise of perturbation-MPRA for predicting non-coding regulatory activities.

## INTRODUCTION

Advances in high-throughput technologies have allowed a detailed characterization of the human genome, including regulatory elements such as enhancers which contain binding motifs for transcription factors (TFs) and play a central role in the transcriptional regulation of gene expression. Aberrations in the non-coding regions of the genome have been linked to numerous polygenic disorders such as cancer, heart, and neurological disorders (1, 2, 3), making the study of non-coding regions an important area of research.

However, linking the non-coding genome to the etiology of diseases is largely limited by the low throughput of conventional “luciferase reporter assays”, especially when numerous non-coding regions are of interest. To address this challenge, massively parallel reporter assays (MPRAs) were developed to simultaneously measure the activity of thousands of regulatory elements and their variants in a single experiment (4, 5). Furthermore, a perturbation-based MPRA approach was introduced to elucidate the regulatory effects of transcription factor (TF) binding motifs, instead of single nucleotide variants (6, 7, 8). The essence of this technique is to analyze the change in the transcription activity of reporter genes after altering the DNA sequence of putative functional regulatory regions.

In our recent studies, we have utilized utilized the perturbation MPRA technique to successfully identify over 500 non-coding genomic regions that temporally regulate gene transcription during neural differentiation (9, 10). Although the potential of perturbation MPRA has been widely acknowledged, there have been limited attempts to evaluate different design strategies of the tested sequences.

Motivated by the scarcity of the gold standard for DNA sequence designing strategies for the MPRAs technique, we propose a framework for assessing and comparing perturbation strategies (Figure 1). Under this framework, we benchmark three different perturbation methods using a publicly available dataset we recently generated (9, 10).

**Figure 1.**
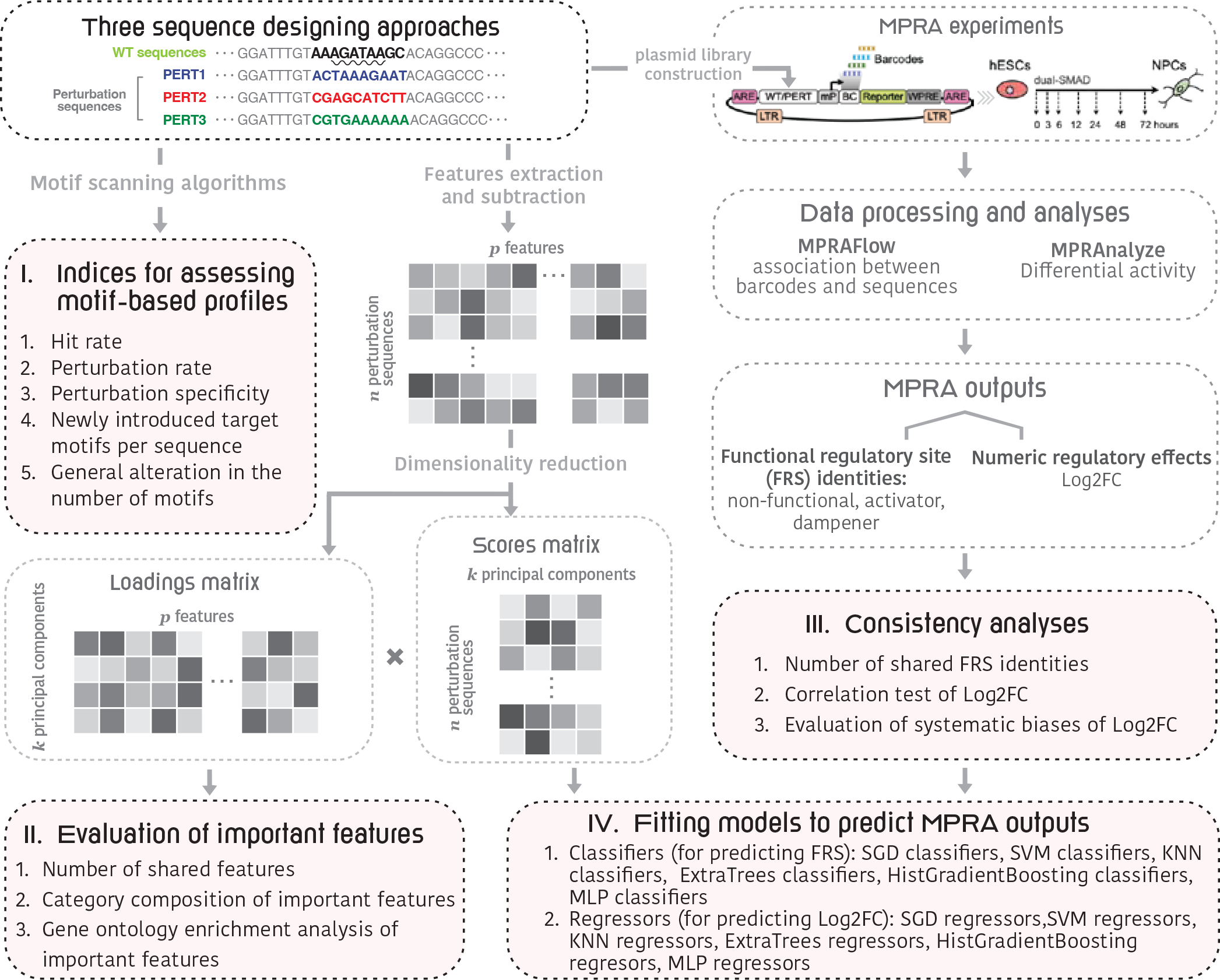
An outline of the framework for evaluation of perturbation-bases massively parallel assays technique. In the “Three sequence designing approaches” box, we used the “GATA known9” motif as an example. In detail, the GATA motifs are a group of sequences conforming to the consensus WGATAR (W = A or T and R = A or G) (marked by the wavy underline), that can be recognized and bound by GATAbinding transcription factors (35).

Briefly, this dataset includes 591 wild-type (WT) sequences, 2,146 motif perturbation sequences, with each sequence perturbed using three different perturbation approaches, and 591 negative control sequences. The perturbation methods, in short, either replaced the target motifs with two different “non-motif” sequences (PERT1 and PERT2), or simply shuffled the nucleotides of target motifs (PERT3).

For benchmarking, we first define five indices to comprehensively evaluate the achievement of the perturbation goals. These indices include, for example, the perturbation rate that indicates the impact on the target motifs both *in-situ* and *ex-situ*, and the specificity index that indicates the proportion of WT motifs that survives the perturbation processes, etc. By comparing these indices, we found that the PERT3 exhibits the highest specificity while the lowest perturbation rate. Next, we compared the consistency of MPRA outputs, both in functional regulatory site (FRS) identities and numeric regulatory effects. Our analyses revealed a high correlation among the three perturbation methods, but we also found a constant bias in the results of PERT1 and PERT2. This is likely due to their insertion of fixed sequences, which may introduce systematic biases to the assayed regions. Finally, we extracted multiple genomic features for each tested sequence and used the difference in the features between the perturbation sequences and their WT equivalents as independent variables to fit predictive machine-learning models. Our results for these predictive models demonstrated the robustness of both classifiers and regressors based on PERT3 data.

To the best of our knowledge, this is the first study that assesses and compares different perturbation methods of MPRA experiments. Our study fills this gap by constructing a blueprint evaluation framework for perturbation sequence designing strategies. Additionally, our results provide guidance for establishing a gold standard of perturbation MPRAs techniques, and our prediction pipeline holds great promise for further computationally identifying functional genomic regulatory regions.

## MATERIALS AND METHODS

### Dataset overview

We utilized a publicly available dataset of perturbation MPRA we recently published (9). The MPRA experiment was performed in the human embryonic stem cell line across seven time points after neural differentiation induction (0, 3, 6, 12, 24, 48, and 72 hours). Specifically, it assayed three groups of genomic sequences: **a)** Wild type group: 591 wild-type sequences (denoted as “WT”): each WT sequence represents a 171-nucleotide genomic region whose regulatory activity differs over time (10), **b)** Motif perturbation group: 2,146 sequences, each containing a single-perturbed motif within the genomic region of its WT equivalent. And each sequence is perturbed using three different perturbation approaches (denoted as “motif PERT1”, “motif PERT2” and “motif PERT3”):

1. PERT1: a motif is replaced with the prefix of an artificially scrambled motif, while using three bp downstream and upstream of the motif in the WT sequence. Under this strategy, the sequence of the perturbed motif is original sequence start”scrambled motif1 prefix”original sequence end.
2. PERT2: similar to PERT1, a motif is replaced with the prefix of another artificially scrambled motif, while keeping the WT starting and ending sequences. Under this strategy, the sequence of the perturbed motif is original sequence start”scrambled motif2 prefix”original sequence end.
3. PERT3: the motif is scrambled by randomly shuffling its nucleotides.

and **c)** Negative control group 1: 591 scrambled sequences (denoted as “SCRAM”). Scrambled sequences are based on WT sequences with shuffled nucleotides, creating a set of negative controls, **d)** Negative control group 2: these are a set of all the 591 WT sequences where we perturbed a sub-sequence in the length of the average motif (12 bp) in a random location within the WT sequence using the same three perturbation methods (denoted as “non-motif PERT1”,”non-motif PERT2”, and “non-motif PERT3”). The non-motifs and motifs are perturbed using the same three perturbation approaches.

The experimental read-out of the perturbed sequences is then subjected to the MPRAnalyze (11) and MPRAflow (12) tools to assess the motif regulatory effect over time, which is represented by the Log2 fold changes (Log2FC) of PERT read-outs compared to the WT and SCRAM at each of the seven time points. The sequences are further classified into two according to the Log2FC values: activating (Log2FC *>* 0) and repressing (Log2FC *<* 0).

Additionally, to identify the functional regulatory sites (FRS), we used MPRAnalyze (11) to apply a set of four filters to the PERT sequences (9):

1. At one or more time points, the activity of a PERT sequence significantly deviates from its WT equivalent.
2. The temporal activity of a PERT sequence significantly deviates from its WT equivalent.
3. The activity of either a PERT sequence (at one or more time points) or a WT sequence (across all the time points) is significantly higher than its corresponding SCRAM negative control sequence.
4. The temporal activity of either a PERT or a WT sequence is significantly higher than its corresponding SCRAM negative control sequence.

The target motif of a sequence will be labeled as an FRS if the sequence passes all four filters and shares consistent effects (either activating or regressing) in PERT3 and either PERT1 or PERT2. In summary, the MPRA output consists of the numeric regulatory effect (Log2FC) and the multi-class FRS identities at seven time points. These two output types are used as input variables for training the prediction models.

### Metrics for assessing motif-based profiles

*Hit rate (HR)* A “hit” sequence indicates the *in-situ* removal of its target motif (*in-situ* removal = “genomic location-specific removal”). In detail, we define “hit” as the target motif of a perturbed sequence that does not occur in the scanning results of the Find Individual Motif Occurrences (FIMO) tool (13), matched by the motif name, DNA strands, and genomic coordinates; otherwise, it’s a “fail.” The hit rate of PERT_i_ is denoted as HR_i_:

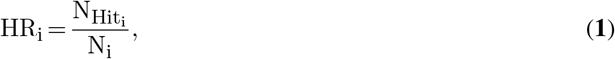

where 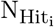 is the number of “hit” sequences and N_i_ is the total number of designed sequences in PERT_i_.

#### Perturbation rate (PR)

A “perturbed” sequence indicates that all motifs that match the target motif ID are removed within the designed genomic region. In detail, we define a sequence as “perturbed” if the name of the target motif occurs in its FIMO scanning results, regardless of its genomic position. Then, the perturbation rate of PERT_i_ is formulated as:

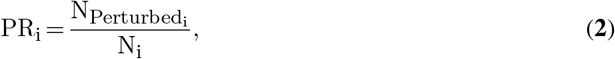

where 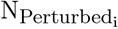 is the number of “perturbed” sequences and Ni is the total number of designed sequences in PERT_i_.

#### Perturbation specificity (PS)

To assess how many WT motifs are impacted by the perturbation, we introduce the “perturbation specificity” metric. For the designed sequence *j* of PERT_i_, its perturbation specificity is formulated as:

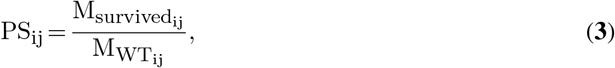

where 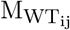 is the number of motifs that overlap with the target motif in the corresponding WT sequence of designed sequence *j* of PERT_i_, and 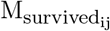 is the occurrence of wild-type motifs that are still present within the designed sequence *j* of PERT_i_. Both 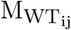 and 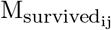 are obtained from FIMO scanning results.

#### Newly introduced target motifs per sequence (NTM)

Since the perturbation process alters the orders of nucleotides, some of the newly introduced motifs may be identical to the target motifs. To assess such impact of the perturbation methods, we calculated and compared the “number of newly introduced target motifs per sequence” among the three perturbation methods. For PERT_i_, its “newly introduced target motifs per sequence” metric is formulated as:

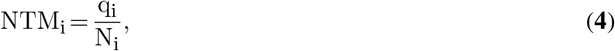

where q_i_ is the number of newly introduced motifs that are identical to the target motif IDs in PERT_i_, and N_i_ is the total number of designed sequences in PERT_i_.

#### General alteration in the number of motifs

To assess the non-specific impacts of the perturbation, we obtained and compared these indices among the three perturbation methods:

1. The number of gained motifs
2. The number of lost motifs
3. The net change in the number of motifs

#### Consistency analysis of MPRA outputs

The MPRA outputs consist of two parts: the multi-labeled FRS identities and the numerical regulatory effects. To analyze the consistency of FRS identities, we counted the number of overlapped and unique activatorsregressors that are specific to their genomic coordinates and DNA strands across three perturbation methods. And the results are visualized by an UpSet plot (14). As for the agreement in numerical regulatory effects, we tested the correlation of Log2FCs between any two of the three perturbation methods using three correlation tests: Pearson *r* correlation, Spearman’s rank correlation, and Kendall’s rank correlation test.

### Features extraction for designed sequences

The features are a major determinant of the performance of predictive models (15, 16). The features used in this work can be grouped into two main categories: sequence-based features and time-specific features.

*Group A sequence-based features*

Since this group of features is based on the nucleotide sequences, each assayed sequence, either WT or perturbed, has its own set of features:

- DNA 5-mer frequencies: 1,024 features indicating the counts of all possible nucleotide 5-mers.
- #5-mers: a single feature summarizing the number of distinct 5-mers.
- DeepBind scores: 515 predicted scores of all pre-trained DeepBind models for transcription factor (TF) binding (17).
- #DeepBind-top: a single feature summarizing the number of models above the 90^th^ percentile across all the DeepBind models for TF binding (17).
- DeepSEA scores: 21,907 chromatin profiles (transcription factor, histone marks, and chromatin accessibility profiles across a wide range of cell types) from the underlying DeepSEA learning model (16).
- #DeepSea-top: a single feature summarizing the number of chromatin profiles above the 90^th^ percentile across all the DeepSEA profiles (16).
- DNA shape metrics: 13 predicted DNA shape features, which are: helix twist (HelT), Rise, Roll, Shift, Slide, Tilt, Buckle, Opening, propeller twist (ProT), Shear, Stagger, Stretch, and minor groove width (MGW) (18, 19).
- Max polyA/polyT lengths: two features indicating the length of the longest polyA and polyT subsequences, respectively.
- #ENCODE/CIS-BP motifs: 4,706 features, showing the number of significant DNA-binding ENCODE/CIS-BP (20, 21, 22) motifs from simple DNA-binding motif scoring using the Find Individual Motif Occurrences (FIMO) tool (13).
- ENCODE/CIS-BP motif summaries: four features indicating the number of motifs, and the maximum number of ENCODE/CIS-BP motifs within a 20 bp window in the sequence, as determined by FIMO scanning algorithm (13, 21, 22).
- #TF family: fourteen features indicating the frequency of major TF families based on the FIMO results of ENCODE/CIS-BP scannings, which are: Basic Domain Group, Beta-Scaffold Factors, Helix-turn-helix, Other Alpha-Helix Group, Unclassified Structure, and Zinc-Coordinating Group (23).

For each perturbed sequence, we subtract its sequence-specific features from that of its WT equivalent. Additionally, we calculate the Levenshtein similarity scores between the perturbed sequences and their respective correspondent WT sequences (24, 25). In total, 28,189 features are yielded from group A.

These differences in features (denoted as ∆”[feature name]”, e.g., ∆#5-mers), along with the Levenshtein similarity scores, are then subject to the feature normalization process (see Section “Feature normalization”).

*Group B: time-specific features*

The time-specific features used in this study are the experimental read-outs of WT sequences (10). These features include the signals of three genomic assays at seven time points (0, 3, 6, 12, 24, 48, and 72 hours):

- ATAC-seq: the normalized number of reads using DESeq2 (26) from an overlapping ATAC-seq peaks within the designed genomic region
- H3K27ac ChIP-seq, the normalized number of reads using DESeq2 (26) from an overlapping H3K27ac peaks within the designed genomic region
- RNA-seq: mRNA expression of the nearest gene to the designed region

In total, three features are yielded from group B. For each perturbed sequence, we use the time-specific feature of its corresponding WT sequence as its feature to fit prediction models.

### Feature normalization

Performing principal component analysis (PCA) is a common technique to reduce the number of features in high-dimensional data to avoid over-fitting and improve the generalization performance of machine learning models. In this case, PCA was applied to the large number of group A features (28,189) to reduce them into a smaller set of principal components (PCs) that capture the maximum amount of variability in the data. By selecting the number of PCs such that they explain at least 99% of the variance in the data, the most important information in the original features is retained while reducing their dimensionality.

In this study, we employed PCA to transform the 28,189 group A features into 1,500 PCs for each perturbation method. Together with the time-specific features of group 2, a total of 1,503 features were used as input for subsequent prediction tasks. This approach helps to prevent over-fitting and improves the accuracy of the machine learning models.

### Calculation of the feature importance scores

We first defined the importance score *I* of feature *i* as the largest loading score of feature *i* across 1,500 PCs. In particular, from the PCA step, we obtain a matrix *L* to denote the loadings matrix that explains the correlations between the original features and the PCs. *L* is a 28,189 × 1,500 matrix with rows representing features and columns representing 1,500 PCs. For feature *i*, its loading score on the *j*^*th*^ dimension is denoted as *L*_*ij*_. We then define the importance score *I* of the feature *i* as its largest loading score across the 1,500 PCs:

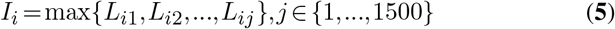

### Gene ontology analysis

We conducted the Gene ontology (GO) over-representation analysis using the genes corresponding to the top 2,500 important TF binding features. The results were determined using the R package ClusterProfiler (27). The significance of GO terms was defined as an FDR-adjusted p *<* 0.05.

### Model training

#### Classification models

We utilized six classification models to predict the FRS identity of perturbed sequences:

1. SGD: linear SVM classifiers with stochastic gradient descent (SGD) training (28)
2. SVC: C-Support vector classifiers (29)
3. KNN: classifiers based on k-nearest neighbors voting (30)
4. ET: ExtraTrees classifiers (31)
5. HGB: histogram-based gradient boosting classifiers (32)
6. MLP: multilayer perceptron classifiers (33)

All classifiers were run with the default settings of the scikit-learn package (34). The 1,503 normalized feature values were used as input. To generate target values, the FRS identity labels at seven time points were concatenated and stacked into a single variable.

#### Regression models

1. SGD: SGD: linear regressors fitted by minimizing a regularized empirical loss with SGD training (28)
2. SVC: SVR: Epsilon-Support vector regressors (29)
3. KNN: regressors based on k-nearest neighbors voting (30)
4. ET: ExtraTrees regressors (31)
5. HGB: histogram-based gradient boosting regressors (32)
6. MLP: multilayer perceptron regressors (33)

All regressors were run with the default settings of the scikit-learn package (34). The 1,503 normalized feature values were used as input. The Log2FCs at seven time points were concatenated and stacked into a single variable, and regarded as target values.

### The randomized 10-fold cross-validation test

We performed 10-fold cross-validation tests to evaluate the performances of different models. A 10-fold cross-validation test was chosen as it provides a good balance between minimizing bias and reducing variance. In detail, the dataset is randomly partitioned into ten subsets, with one subset utilized as the testing dataset and the other nine together as the training data set. This procedure was conducted 10 times, with each subset being used once as a testing dataset to generate ten models. The average performance of these ten models was used to evaluate the performance of the different models.

To ensure a fair and objective comparison among the models, we strictly implemented their algorithms and optimized parameters to build models on the same training dataset and subsequently benchmark their performance on the independent test datasets.

### Model performance measures

The performance of classification models is evaluated using the area under the receiver-operating characteristic curve (AUROC). For the regression models, we evaluated their performance using three correlation tests: Pearson, Spearman, and Kendall. Specifically, we tested the correlation between the predicted Log2FC values and the observed Log2FC values for each fold.

### Statistical tests

For the motif-based profile metrics, the Kruskal–Wallis one-way analysis of variance and post-hoc pairwise Dunn’s multiple comparisons test were used to identify statistically significant differences in continuous variables, including the perturbation specificity and the number of gained/lost motifs. Moreover, the pairwise Fisher’s exact test was conducted to compare the count data, including hit and perturbation rates. The pairwise exact binomial test was performed to compare newly introduced target motifs per sequence (NTM).

For the consistency analyses, the correlation of Log2FCs was indicated by three correlation coefficients: Pearson’s *r*, Spearman’s *ρ*, and Kendall’s *τ* coefficient. The *P* values of correlation tests were subsequently adjusted for multiple comparisons at seven different time points by the Benjamini-Hochberg method.

For the performance evaluation of prediction models, we performed pairwise Wilcoxon rank sum tests on the AUROC and correlation coefficients. For all pairwise tests, a threshold of 0.05 was applied to the *P* values adjusted by the Benjamini-Hochberg method. And an *α* level was considered 0.05 for all statistical tests in this study.

## RESULTS

To evaluate the three perturbation methods, we first defined five motif-based metrics: *1) hit rate*, representing the rate of *in-situ* motif perturbation, defined as the proportion of designed sequences that successfully eliminate the target motif at the target genomic locale, *2) perturbation rate*, which represents the rate of both *ex-situ* and *in-situ* motif perturbation and is defined as the proportion of designed sequences that eliminate all the motifs that match the target motif ID within the 171-nucleotide genomic region, *3) perturbation specificity*, indicating the global impact of perturbation on all the motifs that lie within the perturbed sequence, and is defined as the proportion of WT motifs that are still found in the perturbation sequence, *4) newly introduced target motifs per sequence*, which reflects the occurrence of gained motifs that are identical to the target motif ID, and is calculated by dividing the total number of such gained motifs by the total number of perturbation sequences, *5) non-specific changes in the number of motifs*, which include the number of gained, lost motifs, as well as the net change in the number of motifs within the perturbation sequence. We then assess the differences in these metrics across the three perturbation methods (Figure 1, part I). Next, we assess the important features representing variability among all perturbation methods (Figure 1, part II). Third, we compare the consistency of MPRA outputs (Figure 1, part III). Finally, to evaluate the generalizability in referencing the non-coding regulatory activity of the three perturbation methods, we compare how different prediction models perform across the three perturbation methods (Figure 1, part IV).

### All perturbation methods achieve high *in-situ* hit rates

The basic goal of a motif perturbation is to remove the target motif at the target genomic location. To assess how well each perturbation method is in reaching this goal, we computationally identified the occurrences of the motifs in perturbed sequences, by using the FIMO tool 13 scanning results and matching the motif names, DNA strands, and genomic coordinates (Section “Methods”).

If a perturbed sequence yields a “non-occurrent” result, it is defined as a “hit” indicating a successful perturbation, otherwise a “fail” (Section “Methods”; Figure 2A). We then calculated and compared the proportion of hit and fail sequences for each perturbation method (equation **1**). The hit rates of PERT1 and PERT2 are similar (HR_1_ = 98%, HR_2_ = 99%), and both are significantly higher than that of PERT3 (HR_3_ = 98%, pairwise Fisher’s exact test, PERT1 vs. PERT2, *P* = 1.00; PERT1 vs. PERT3, *P* = 1.42×10^*−*3^; PERT2 vs. PERT3, *P* = 1.42 × 10^*−*3^). Still, all three PERTs exhibit high hit rates of over 98% (Figure 2B).

**Figure 2.**
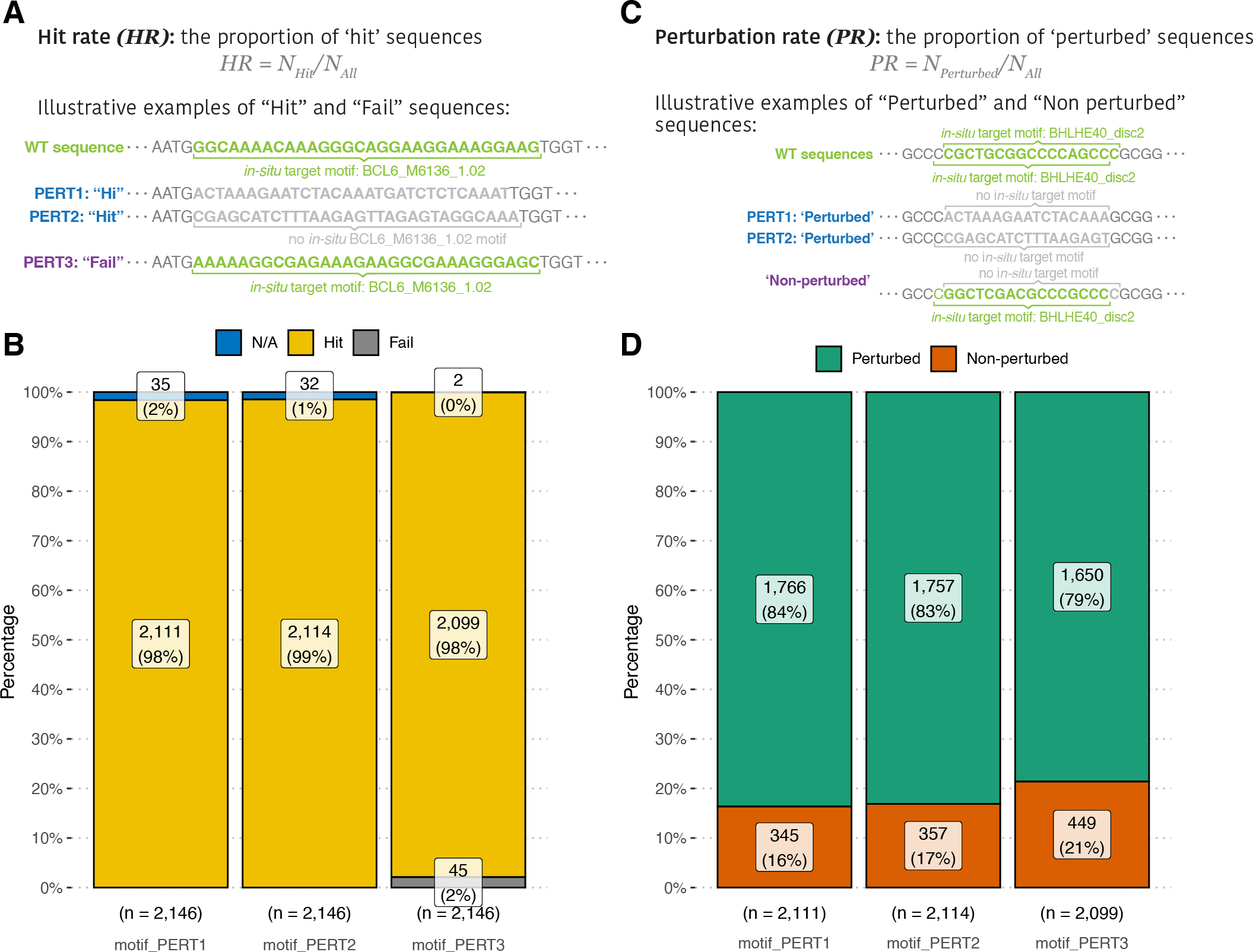
Evaluations of perturbation-wise metrics. **(A)** Toy examples of “hit” and “fail” sequences. **(B)** A comparison of hit rates among three perturbation methods. **(C)** Toy examples of “perturbed” and “non-perturbed” sequences. **(D)** A comparison of perturbation rates among three perturbation methods.

### The non-location-specific perturbation rate of PERT3 is the lowest

Apart from the basic goal, one of the advanced goals of motif perturbation is to reduce the regulatory activity of the target motif to the baseline, that is, to eliminate all the motifs that are identical to the target motif ID within the 171-nucleotide genomic region of perturbation sequence. Hence, we further quantified the occurrence of the target motif in each “hit” sequence using the FIMO scanning results, by matching only the motif name and not its location. Sequences were defined as “perturbed” if no designed target motif was found within their genomic region, and the perturbation rate was then calculated as the proportion of ‘perturbed’ sequences (Figure 2C). In simple words, this metric indicates the rate of both *ex-situ* and *in-situ* motif perturbation, that is not specific to the target genomic location (Section “Methods”, equation **2**).

Comparing the perturbation rate of the three PERTs, we found that PERT1 and PERT2 possess similar perturbation rates of over 80%. Although the perturbation rate of PERT3 is significantly lower than those of the other two, it is still as high as 79% (Figure 2D, *PR*^1^ = 84%, *PR*^2^ = 83%, *PR*^3^ = 79%; pairwise Fisher’s exact test, PERT1 vs. PERT2, *P* = 0.649; PERT1 vs. PERT3, *P* = 8.79 × 10^*−*5^; PERT2 vs. PERT3, *P* = 4.58 × 10^*−*4^). These results indicate that the strategic design of perturbation sequences (PERT1 and PERT2), instead of simply shuffling the nucleotide sequences (PERT3), leads to a higher chance of perturbing non-location-specific target motifs within genomic regions.

### Perturbation specificity are similar among three methods

Another advanced goal of motif perturbation is to keep the impact on the overall motifs as low as possible: since the perturbation process essentially alters the DNA sequence within a certain range of the genome, the motifs that overlap with the target motifs are likely to be affected. To assess such a global impact of the perturbation on all the motifs that lie within the perturbation sequence, we introduced the perturbation specificity metric. It is defined as “the proportion of WT motifs that are still present within the genomic region after perturbation” (Section “Methods”, equation **3**, Figure 3). Comparing the perturbation specificity among three PERTs, we found that all three perturbation methods vastly affect the WT motifs. Namely, only 10% of the overlapping WT motifs “survived” the perturbation processes. Specifically, PERT3 has the highest perturbation specificity, which implies that randomly shuffling nucleotides exerts the least overall impact within the genomic regions of perturbed sequences (Figure 3B, *PS*^1^ = 7%, *PS*^2^ = 7%, *PS*^3^ = 11%; pairwise Dunn’s test, PERT1 vs. PERT2, *P* = 5.73 × 10^*−*3^; PERT1 vs. PERT3, *P* = 8.95 × 10^*−*27^; PERT2 vs. PERT3, *P* = 1.14 × 10^*−*15^).

**Figure 3.**
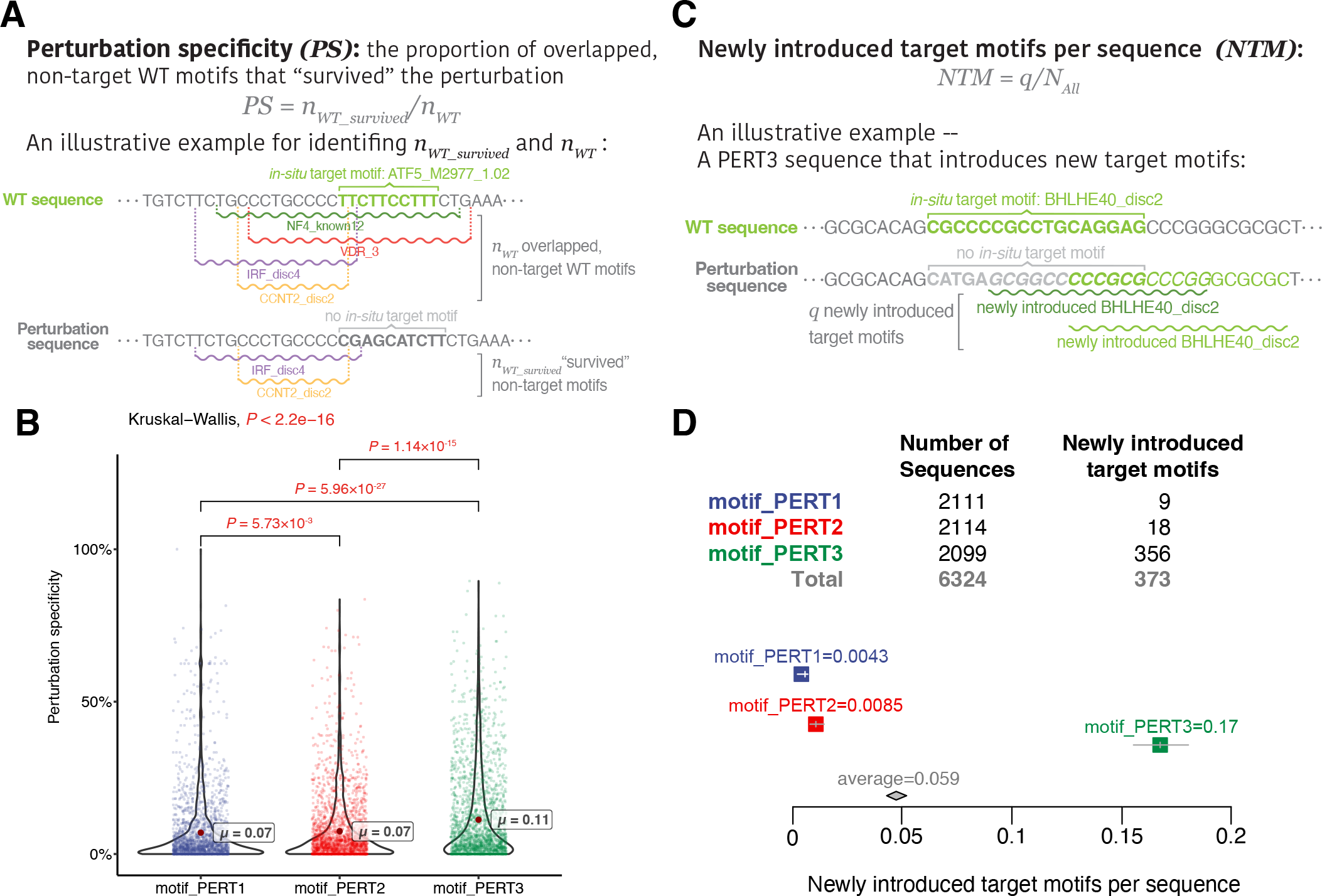
Evaluations of motif-based metrics. **(A)** An toy example of calculating perturbation specificity. **(B)** A comparison of perturbation specificity among three perturbation methods. Significant *P* values (*P* < 0.05) are shown in red. **(C)** An toy of calculating “newly introduced target motifs per sequence”. **(D)** A comparison of “newly introduced target motifs per sequence” among three perturbation methods.

On the other hand, another advanced goal is to avoid “creating” target motifs in the perturbation sequences. To this end, we sought to investigate which perturbation approach introduces the highest number of new motifs that are identical to the target motif ID. We defined the newly introduced target motifs per sequence metric, which is calculated by dividing the total number of “newly introduced target motifs” by the total number of sequences for each perturbation method (Section “Methods”, equation **4**, Figure 3C). The highest metric is produced by PERT3, indicating that shuffling the nucleotides increases the probability of generating the same motifs as the target ones (Figure 3D, *NTM*^1^ = 0.0043, *NTM*^2^ = 0.0085, *NTM*^2^ = 0.17; pairwise exact binomial test, PERT1 vs. PERT2, *P* = 0.122; PERT1 vs. PERT3, *P* = 2.35 × 10^*−*92^; PERT2 vs. PERT3, *P* = 1.73 × 10^*−*82^).

### All three perturbation approaches vary in motif gain/loss

To gain a better perturbation effect, the impacts that are non-specific to the target motifs should also be minimized as much as possible. To address such impacts, we evaluated the overall motifs gained or lost across motif perturbation approaches (Figure 4A), and found that PERT3 gains significantly over 30 more motifs than PERT1 and PERT2 (Figure 4B, *PERT* 1∼ = 8.45, *PERT* 2∼ = 10.63, *PERT* 3∼ = 43,26; pairwise Dunn’s test, PERT1 vs. PERT2, *P* = 1.18 × 10^*−*5^; PERT1 vs. PERT3, *P* = 1.55 × 10^*−*206^; PERT2 vs. PERT3, *P* = 2.18 × 10^*−*199^). However, the number of motifs lost was similar among the three methods (Figure 4C, *PERT* 1∼ = 101.12, *PERT* 2∼ = 99.95, *PERT* 3∼ = 90.73; pairwise Dunn’s test, PERT1 vs. PERT2, *P* = 0.771; PERT1 vs. PERT3, *P* = 0.0811; PERT2 vs. PERT3, *P* = 0.109).

**Figure 4.**
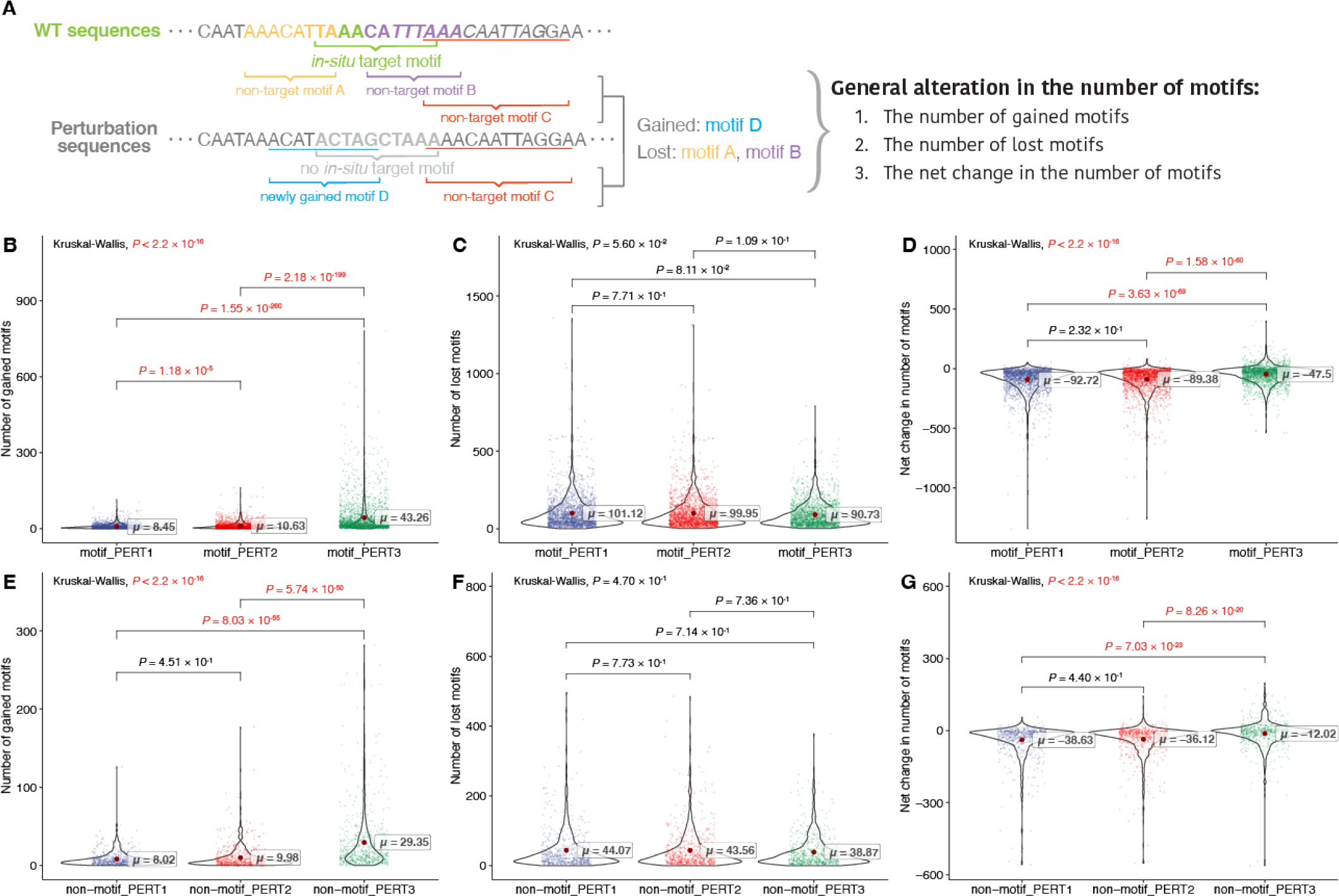
Evaluation of general alteration in the number of motifs. **(A)** Toy examples of calculating general alteration in the number of motifs. **(B-D)** The results for motif perturbations: **(B)** the number of gained motifs, **(B)** the number of lost motifs, and **(B)** the net change in the number of motifs. Significant P values (P *<* 0.05) are shown in red. **(E-G)** The results for non-motif perturbations: **(E)** number of gained motifs, **(F)** number of lost motifs, and **(G)** net change in the number of motifs. Significant P values (P *<* 0.05) are shown in red.

We also compared the net change in the number of motifs for each perturbation approach. We observed that PERT3 resulted in a significantly greater net change compared to the other two approaches, whereas there was no significant difference between PERT1 and PERT2 (Figure 4D, *PERT* 1 ∼=−92.72, *PERT* 2 ∼=−89.38, *PERT* 3 ∼= −47.50; pairwise Dunn’s test, PERT1 vs. PERT2, *P* = 0.232; PERT1 vs. PERT3, *P* = 3.63 × 10^*−*69^; PERT2 vs. PERT3, *P* = 1.58 × 10^*−*60^).

We then compared these non-specific metrics for the non-motif perturbation sequences. We found similar results to the motif perturbation group: PERT3 resulted in the most motif gains (Figure 4E, *PERT* 1∼ = 8.02,*PERT* 2∼ = 9.98,*PERT* 3∼ = 29.35; pairwise Dunn’s test, PERT1 vs. PERT2, *P* = 0.451; PERT1 vs. PERT3, *P* = 8.03 × 10^*−*55^; PERT2 vs. PERT3, *P* = 5.74 × 10^*−*50^), with no significant difference in the number of lost motifs (Figure 4F, *PERT* 1∼ = 44.07,*PERT* 2∼ = 43.56,*PERT* 3 ∼= 38.67). In addition, the net change in the number of motifs of PERT3 is negative but the highest (Figure 4G, *PERT* 1∼ = −38.63,*PERT* 2∼ = −36.12,*PERT* 3∼ = −12.02; pairwise Dunn’s test, PERT1 vs. PERT2, *P* = 0.44; PERT1 vs. PERT3, *P* = 7.03 × 10^*−*23^; PERT2 vs. PERT3, *P* = 8.26 × 10^*−*20^). These findings further support that the differences in the non-specific impacts are due to the perturbation method used.

### The three perturbation approaches share similar important features, specifically neural developmental features

We then set out to investigate which innate features represent the variances among perturbation sequences, and whether these features differ using different perturbation methods. First, we queried the top 10% of the features (2,500) that explain the variability among perturbed sequences (Section “Methods”), and found that a majority of these features (1,601) are shared by at least two perturbation methods (Figure 5A). Notably, these features mainly fall into “the change in the number of ENCODECIS-BP motifs” and “5-mers frequencies” categories.

**Figure 5.**
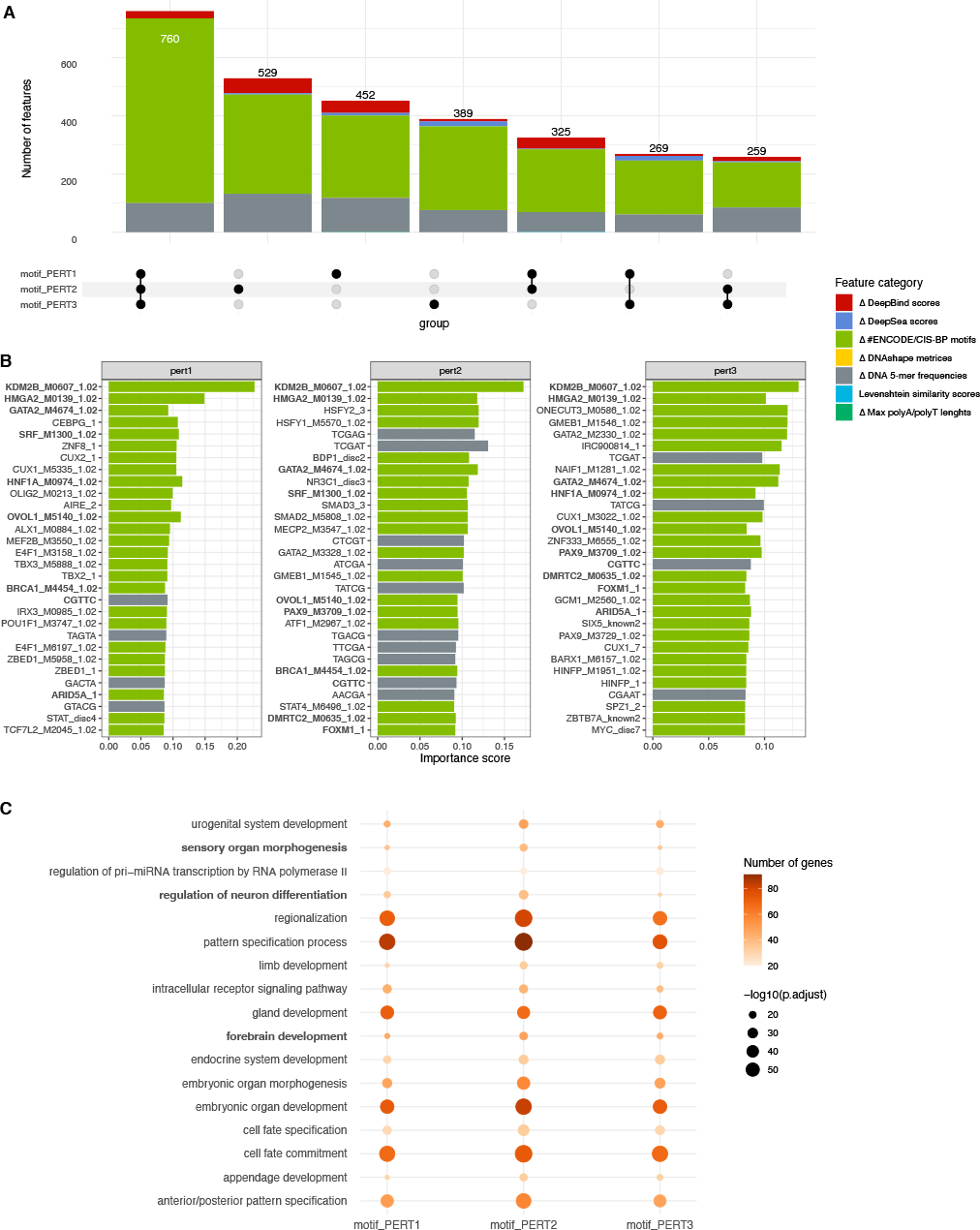
Assessment of the important features representing perturbation sequences. **(A)** The number of important features shared by three perturbation methods. **(B)** Top 30 important features of each perturbation method. The names of features that are shared by at least two perturbation methods are marked in bold. **(C)** Gene ontology enrichment analysis of the top 2500 genes represented by the TF binding factors.

Further scrutiny of the top 30 features revealed a substantially large overlap among the three perturbation methods (Figure 5B). Since a majority of the shared features are transcription factor (TF) binding motifs, we conducted gene ontology analysis on the TFs corresponding to the top 2,500 binding motifs. The analysis revealed consistent enrichment of early embryonic development ontologies, including neural development pathways among three perturbation approaches (Figure 5C). These findings suggest that the three perturbation approaches share important features related to neural development.

### The MPRA outputs are largely consistent across different perturbations

After assessing the basic and advanced goals of perturbation methods, we next evaluated the consistency of MPRA outputs among three perturbation methods. The MPRA output consists of two parts: the multi-class FRS identities, and the numeric regulatory effect (Log2FC) at seven time points of neural differentiation (Section “Methods”).

For the FRS identities, the activities of 419 functional regulatory sites are consistent across three perturbation methods, and 95% (399) of them are activators (Figure 6A). Additionally, 262 sites are consistent in any of the two approaches but not the remaining one (Figure 6A). In terms of the Log2FC, we found a high correlation among all three perturbations across all the time points (Figure 6B-D). However, we found that PERT2 yielded higher Log2FC than the other two approaches (Supplementary Figure 1). This indicates that using a perturbation approach where the same sequence is being introduced, can cause a constant bias in the results (e.g., higher or lower Log2FC for PERT2 or PERT1 respectively).

**Figure 6.**
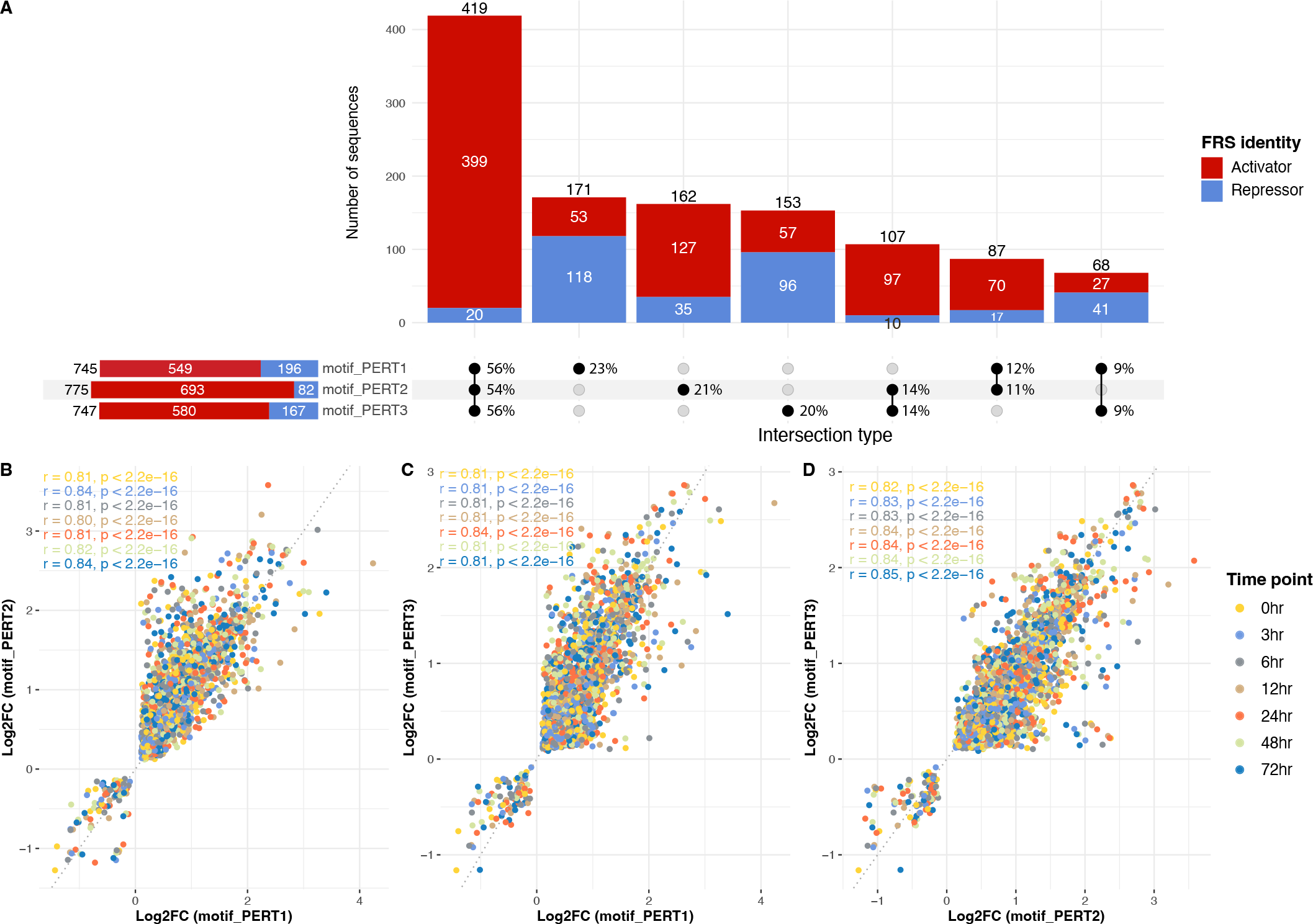
Consistency of MPRA outputs among three perturbations. **(A)** Number of sequences that share the same FRS identities. The bars are colored by activators (red) and repressors (blue). In the “intersection type” matrix. The percentages are row-normalized, indicating the proportion of sequences belonging to different intersection types within each perturbation method. **(B)** The correlation of Log2FC between motif_PERT1 and motif_PERT2. Each dot is a perturbation sequence and is colored by the time point. **(C)** The correlation of Log2FC between motif_PERT1 and motif_PERT3. **(D)** The correlation of Log2FC between motif_PERT2 and motif_PERT3.

### Predictive models of MPRA activity perform the best in PERT3

The perturbation MPRA technique, if designed appropriately, has the potential to predict the activity of non-coding regulatory genomic regions 15. Namely, it is feasible to predict the regulatory activity of a motif by fitting predictive models using the difference in the features between its WT sequence and perturbation sequence. Consequently, this leads to a critical question: which sequence design method for motif perturbation could yield the best performance of such prediction models? This suggests that by designing the perturbation sequences, we may expand the applicability of perturbation MPRA from experimentally identifying regulatory motifs only within designed genomic regions to computationally predicting regulatory elements throughout the non-coding genome. In light of this, we further compared the performances of three perturbation methods using the supervised models as described in the Methods section.

Briefly, we use the difference of features between perturbation sequences and their equivalent WT sequence as the independent variables to fit both classification and regression models. Next, we perform a 10-fold cross-validation for each perturbation data. To benchmark the performance of the models, we statistically compared the AUROC for classifiers and the Pearson correlation coefficient for regressors on the independent test data sets in each fold.

For the classification models that predict the measure of motif FRS identities, we report the receiver-operating characteristic curve (AUROC) of three perturbation approaches (Figure 7). We found that three non-linear models (ET, HGB, and MLP) exhibit high robustness in predicting the FRS identities in the three perturbations. Furthermore, using the results from ET models, we found that PERT3 significantly outperforms PERT2 and PERT1, and PERT1 significantly outperforms PERT2 (pairwise Wilcoxon rank sum test, PERT1 vs. PERT2, *P* = 5.58 × 10^*−*5^; PERT1 vs. PERT3, *P* = 3.24 × 10^*−*5^; PERT2 vs. PERT3, *P* = 3.24 × 10^*−*5^).

**Figure 7.**
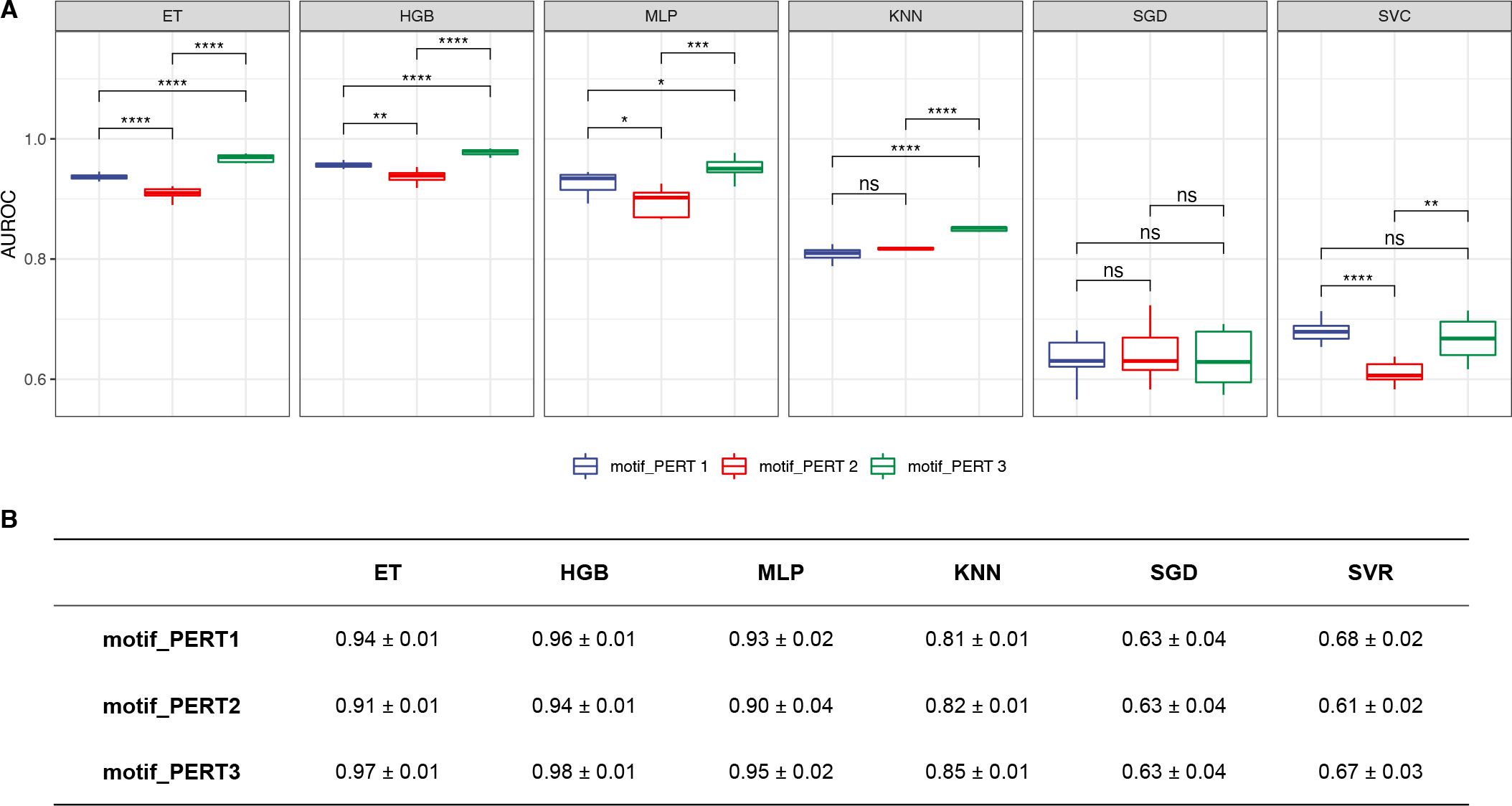
Performance of classification models. **(A)** The area under the receiver-operating characteristic curve (AUROC) of different classification models. Asterisks/ns indicate levels of statistical significance, calculated by pairwise Wilcoxon rank sum tests (P-value *<* 0.05 *, *<* 0.01 **, *<* 0.001 ***, *<* 0.0001 ****; ns, non significant). **(B)** A summary of the mean *±* standard deviation values for AUROCs of classification models.

For the regression models that predict the quantitative measure of motif regulatory effect, we report the Pearson correlation coefficients for the three perturbation approaches (Figure 8, Supplementary Figure 2, Supplementary Figure 3). Similarly, the model-wise comparison shows the robustness of the ET and HGB model, and PERT3 significantly outperforms the other two methods, while PERT2 outperforms PERT1 (pairwise Wilcoxon rank sum tests, PERT1 vs. PERT2, *P* = 2.57×10^*−*3^; PERT1 vs. PERT3, *P* = 3.89×10^*−*5^; PERT2 vs. PERT3, *P* = 3.89×10^*−*5^).

**Figure 8.**
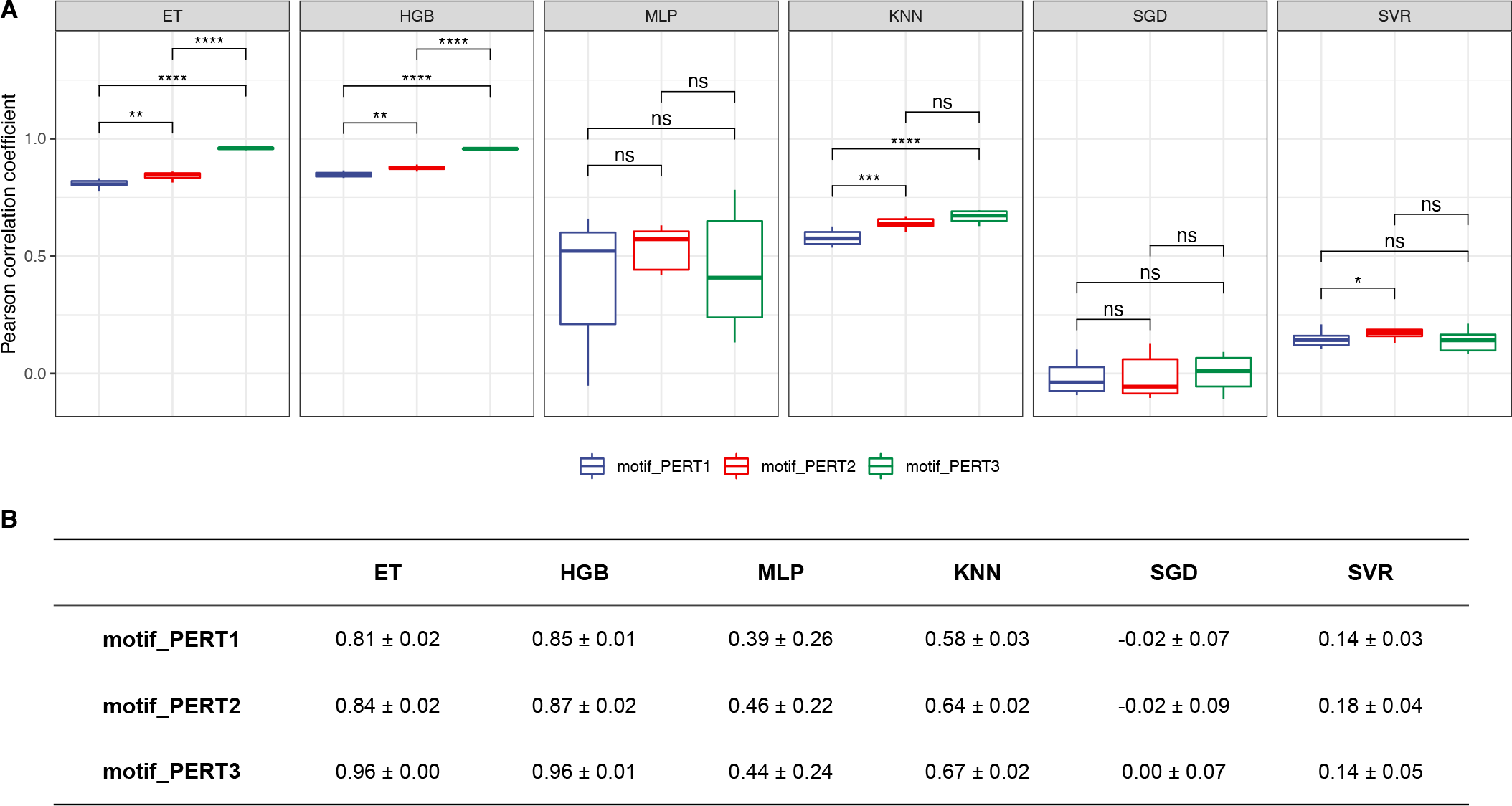
Performance of regression models. **(A)** The Pearson correlation coefficients of different regression models. Asterisks/ns indicate levels of statistical significance, calculated by pairwise Wilcoxon rank sum tests (P-value *<* 0.05 *, *<* 0.01 **, *<* 0.001 ***, *<* 0.0001 ****; ns, non significant). **(B)** A summary of the mean *±* standard deviation values for Pearson correlation coefficients of regression models.

## DISCUSSION

Comprehensively deciphering the regulatory activity of non-coding loci is crucial to the understanding of gene expression dynamics. Shedding light on this, the perturbation-based MPRA technique has enabled the identification of regulatory elements such as enhancers, promoters, and silencers (6, 9, 10). However, insufficient attention has been given to the comprehensive evaluation of various perturbation approaches. As a result, a gold standard of perturbation sequence design strategies remains scant.

Motivated by this scarcity, we proposed a framework for assessing different perturbation approaches, with the aim of better identifying regulatory elements using the perturbation-based MPRA technique. Further, we took advantage of a publicly available data set, which contains the MPRA results acquired from three perturbation approaches (PERT1, PERT2, and PERT3), to conduct an all-inclusive characterization and comparison of these approaches. In short, PERT1 and PERT2 replaced the target motifs with two different “non-motif” sequences, and PERT3 simply shuffled the nucleotides of target motifs.

Starting from the essential ideas of perturbation, which is to eliminate the regulatory effects from target motif(s) within a certain genomic region, we first defined five metrics for assessing the impact from different perturbation approaches (hit rate, perturbation rate, perturbation specificity, newly introduced target motifs per sequence, and general alteration in the number of motifs, see Section “Methods”). These metrics allowed us to scrutinize the overall modification of motif-based profiles within perturbation sequences from different perspectives. Based on our findings, the three approaches exhibit consistently high rates of removing the target motifs at their targeted locations, which indicates success in in-situ motif perturbation. Additionally, the perturbation rate is kept high across the three perturbation methods (80%), with PERT3 being the lowest (79%), while not significantly different. This implies a further achievement in both in-situ and ex-situ removal of target motifs of the three methods. We note that PERT3 shows a higher probability of introducing target-identical motifs. Despite these, PERT3 brings minimal alterations to the WT motifs within the sequence region, implying that the perturbation specificity of PERT3 is the highest. Moreover, PERT3 leads to the least non-specific motif changes. So far, our observation suggests that the selection of perturbation approaches is a trade-off: for the researchers, it becomes a question of whether to sacrifice the perturbation specificity to achieve a high perturbation rate, or whether to pursue a higher specificity at the cost of a lower perturbation rate.

The next part of our framework is the comparison of MPRA outputs since they are crucial for inferring the activity of target motifs. Particularly, MPRA outputs consist of two parts: 1) the functional regulatory site (FRS) identities that indicate whether the target motif is a non-functional, repressing, or activating element, 2,) the numeric regulatory effects (Log2FC) that quantify the FRS motifs. According to our results, the FRS identities are largely consistent and the Log2FC are highly correlated among all three perturbations. Yet, we also observed a constant skew in the results of PERT1 and PERT2, which indicates that inserting repeated/fixed sequences across the assayed regions is likely to introduce systematic biases in downstream results. The results of this part demonstrated that PERT3 is less likely to introduce systematic biases in MPRA outputs, albeit the high-consistency and high-accuracy profiling for the regulatory activity across all three perturbation methods.

The final part of the framework is to evaluate the potential of perturbation-MPRA in predicting the regulatory activity of non-coding motifs, since our previous works have shown robustness in predicting the activity of putative regulatory elements (15, 36). Specifically, by adequately designing perturbation sequences, the MPRA outputs could be computationally predicted by machine-learning models using the biological features of designed sequences as predictor variables. This approach, in some cases, can efficiently identify functional regulatory regions so as to reduce the time and cost of wet lab experiments. Therefore, we developed data-driven models to predict the regulatory activity of target motifs by using the difference in over 28,000 predictive features between perturbation and wild-type sequences. Comparing the performance of models that are built upon the three perturbation methods, we found that PERT3 significantly outperforms the other two in both classification and regression tasks. These findings further support the notion that using a perturbation approach where the nucleotides are being shuffled randomly, works generally better than a replacement with a constant “non-motif” sequence approach.

In summary, we proposed a framework for the evaluation of perturbation sequence design strategies for MPRA experiments, and we utilized this framework to compare three perturbation-based MPRA approaches. From a computational perspective, this study is the first to comprehensively evaluate the library design of the MPRA technique. From an experimental perspective, our results provide deep insights into understanding the impacts of motif perturbation in MPRA experiments. Although it is challenging to offer strict guidance in the absence of *in-vivo* ground truth, we recommend designing sequences by randomly shuffling the nucleotides of the perturbed site when possible.

We anticipate that our findings, together with the proposed framework, will instill a new momentum for the non-coding genomic studies using MPRA techniques, as well as inspire the development of novel comprehensive computational methods. Such efforts and studies will continually contribute to improving our understanding of the functional effects of non-coding regulatory elements.

## ACKNOWLEDGEMENTS

We would like to acknowledge the Office of Advanced Research Computing (OARC) at Rutgers, The State University of New Jersey for providing access to the Amarel cluster and associated research computing resources that have contributed to the results reported here. URL: https://oarc.rutgers.edu

## DATA AND CODE AVAILABILITY

The datasets are available at the NCBI Gene Expression Omnibus (GEO) as accession number GEO: GSE115046.

## SUPPLEMENTARY FIGURES AND TABLES

**Figure Supplementary Figure 1.**
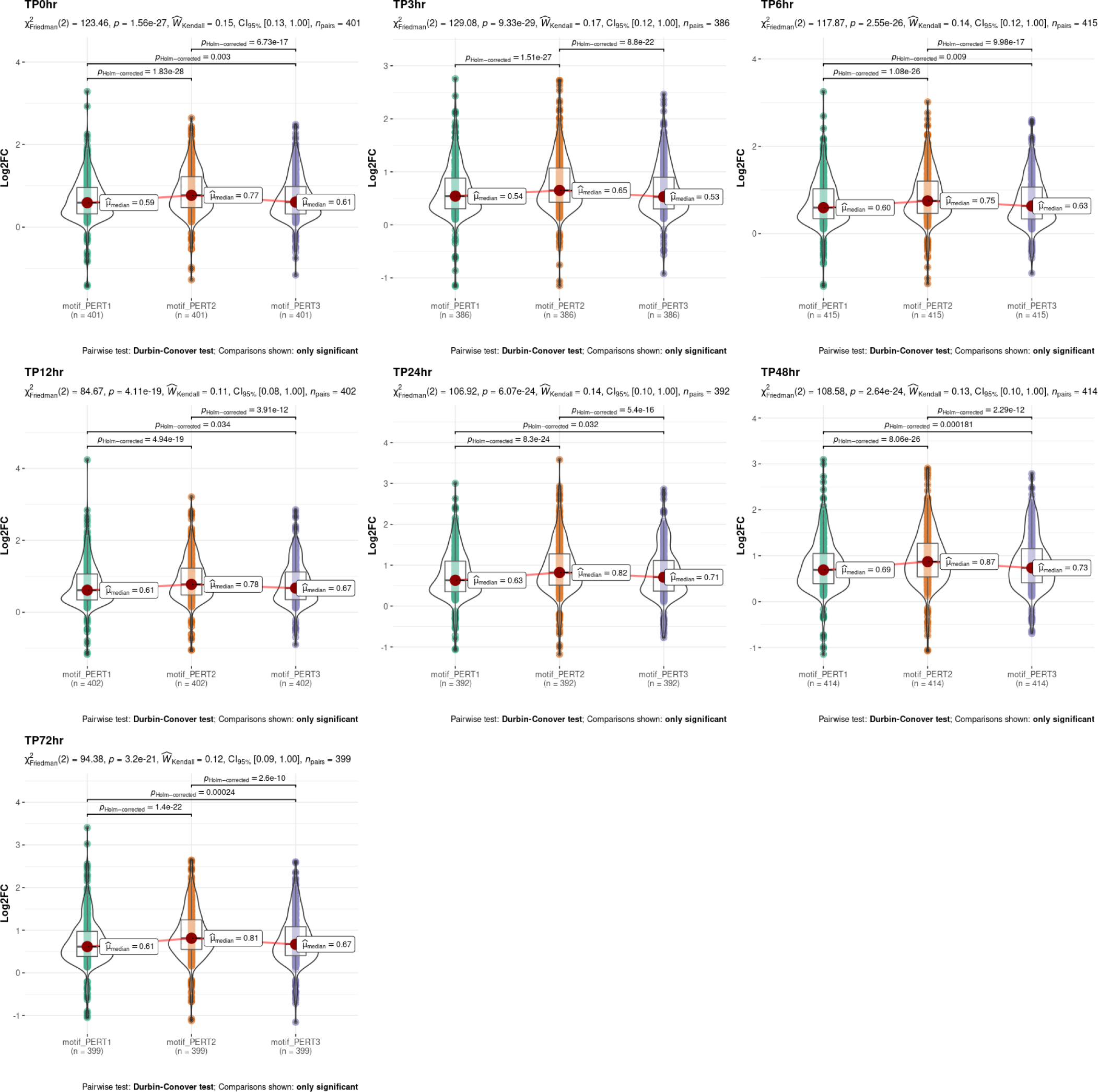
Comparison of Log2FC among three perturbation methods.

**Figure Supplementary Figure 2.**
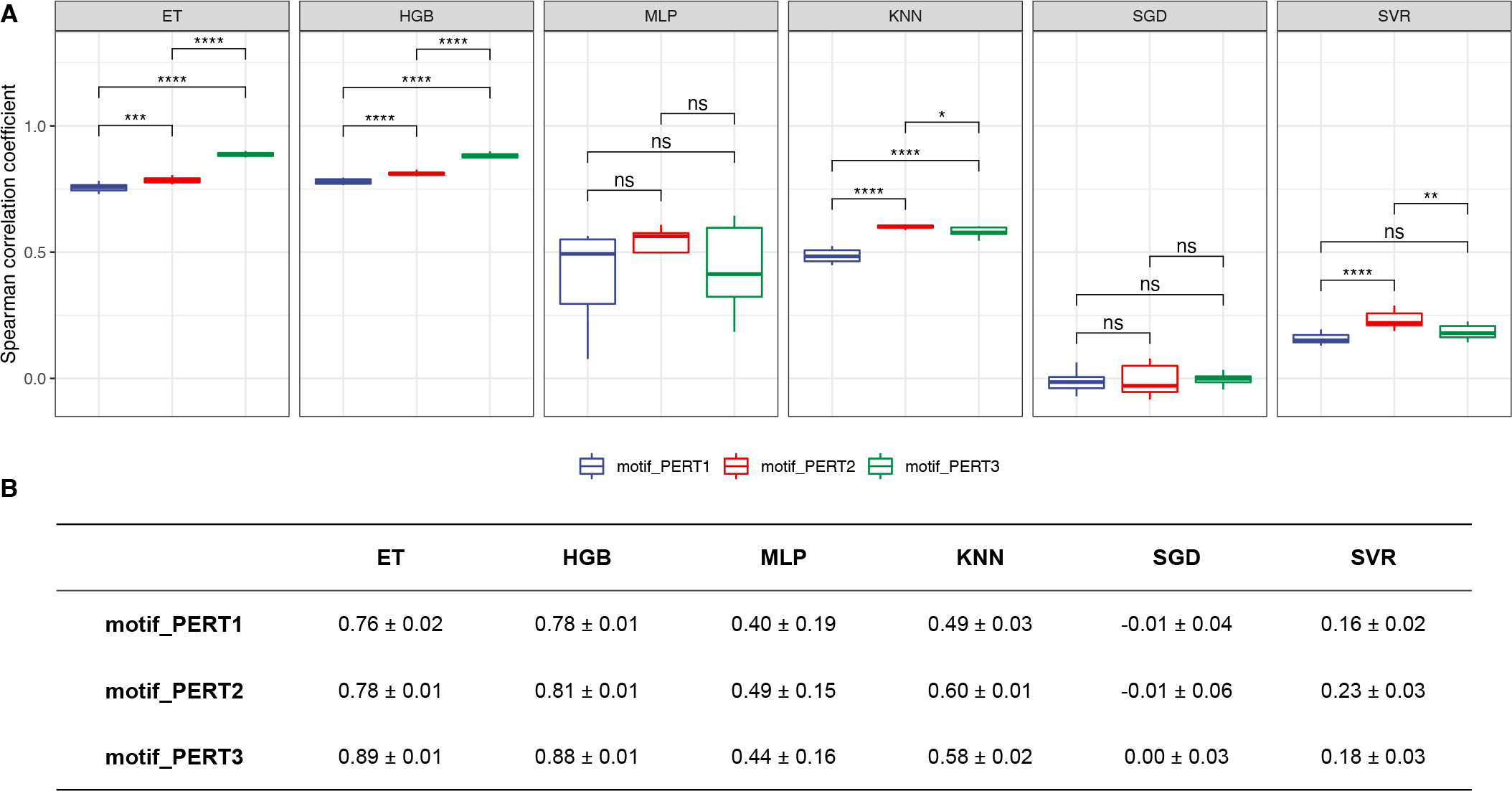
Performance of regression models. **(A)** The Spearman correlation coefficients of different regression models. Asterisks/ns indicate levels of statistical significance, calculated by pairwise Wilcoxon rank sum tests (P-value *<* 0.05 *, *<* 0.01 **, *<* 0.001 ***, *<* 0.0001 ****; ns, non significant). **(B)** A summary of the mean *±* standard deviation values for Spearman correlation coefficients of regression models.

**Figure Supplementary Figure 3.**
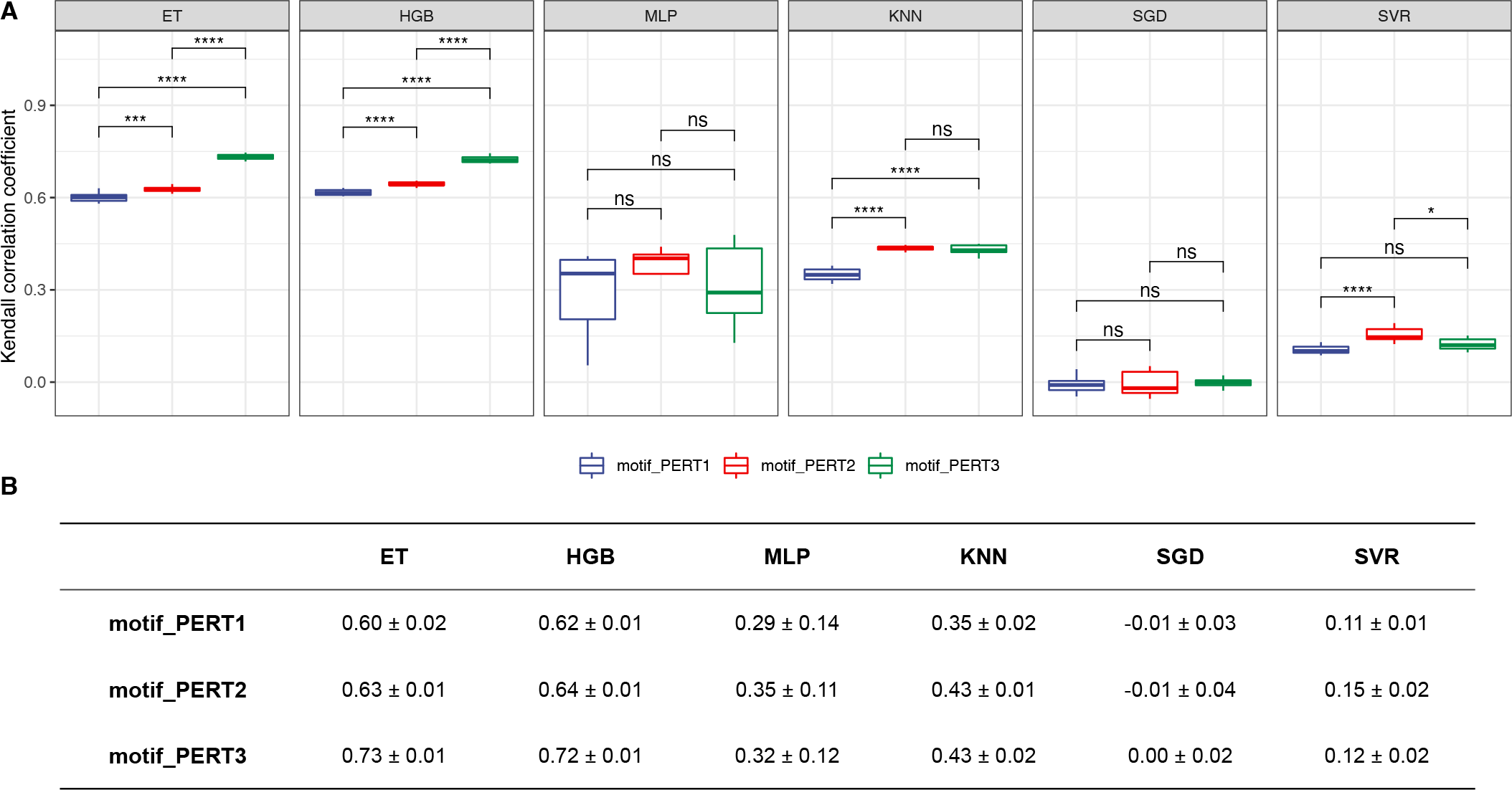
Performance of regression models. **(A)** The Kendall correlation coefficients of different regression models. Asterisks/ns indicate levels of statistical significance, calculated by pairwise Wilcoxon rank sum tests (P-value *<* 0.05 *, *<* 0.01 **, *<* 0.001 ***, *<* 0.0001 ****; ns, non significant). **(B)** A summary of the mean *±* standard deviation values for Kendall correlation coefficients of regression models.

